# SARS-CoV-2 mRNA Vaccine Induces Robust Specific and Cross-reactive IgG and Unequal Strain-specific Neutralizing Antibodies in Naïve and Previously Infected Recipients

**DOI:** 10.1101/2021.06.19.449100

**Authors:** Tara M. Narowski, Kristin Raphel, Lily E. Adams, Jenny Huang, Nadja A. Vielot, Ramesh Jadi, Aravinda M. de Silva, Ralph S. Baric, John E. Lafleur, Lakshmanane Premkumar

## Abstract

With the advance of SARS-CoV-2 vaccines, the outlook for overcoming the global COVID-19 pandemic has improved. However, understanding of immunity and protection offered by the SARS-CoV-2 vaccines against circulating variants of concern (VOC) is rapidly evolving. We investigated the mRNA vaccine-induced antibody responses against the referent WIV04 (Wuhan) strain, circulating variants, and human endemic coronaviruses in 168 naïve and previously infected people at three-time points. Samples were collected prior to vaccination, after the first and after the second doses of one of the two available mRNA-based vaccines. After full vaccination, both naïve and previously infected participants developed comparable robust SARS-CoV-2 specific spike IgG levels, modest IgM and IgA binding antibodies, and varying degrees of HCoV cross-reactive antibodies. However, the strength and frequency of neutralizing antibodies produced in naïve people were significantly lower than in the previously infected group. We also found that 1/3^rd^ of previously infected people had undetectable neutralizing antibodies after the first vaccine dose; 40% of this group developed neutralizing antibodies after the second dose. In all subjects neutralizing antibodies produced against the B.1.351 and P.1 variants were weaker than those produced against the reference and B.1.1.7 strains. Our findings provide support for future booster vaccinations modified to be active against the circulating variants.

## Introduction

The SARS-CoV-2 virus causes a spectrum of disease from asymptomatic to severe forms with high mortality. The nucleocapsid protein encapsulating viral RNA and the surface exposed spike protein are the primary targets of human antibodies. The spike protein of SARS-CoV-2 mediates virus attachment and entry into host cells. It comprises a highly variable S1 segment, which harbors the N-terminal domain (NTD) and the receptor-binding domain (RBD), and a more conserved S2 segment which includes the fusion peptide and heptad repeats required for virus fusion to host cells. Neutralizing antibody response is at present the best correlate of protection^1^. However, the adaptive immune response to SARS-CoV-2 infection is variable^2–4^. RBD accounts for ~90% of the neutralizing activity in SARS-CoV-2 immune sera^5,6^. RBD-specific antibodies target distinct antigenic sites and exert neutralizing activity principally by interfering with spike protein interactions with its cognate receptor, angiotensin-converting enzyme 2 (ACE2). A subset of the NTD-specific antibody also neutralizes SARS-CoV-2 by targeting a supersite, possibly preventing proteolytic activation, membrane fusion, or spike protein interactions with an auxiliary receptor^7^. The spike protein has been targeted in most SARS-CoV-2 vaccines under development and in those approved and currently being administered worldwide.

The recent effort to achieve widespread vaccination against SARS-CoV-2 has left in its wake a host of questions about whether the vaccine can protect against SARS-CoV-2 infection and whether the vaccine can boost immunity in previously infected people ^8–10^. Infection with four human endemic coronaviruses (HCoVs; OC43, HKU-1, NL63, and 229E) are quite common, and most adults have antibodies to these viruses^8,9^. Induction of cross-reactive HCoV antibodies has been reported in SARS-CoV-2 infection, and after vaccination^11^. While their role in protection or immunopathogenesis remains unclear^12,13^, levels of HCoV cross-reactive antibodies correlate with disease severity^14^. The emergence of new, increasingly infectious and virulent SARS-CoV-2 variants is causing significant concern in global human health. The US SARS-CoV-2 Interagency Group has classified the variants currently circulating in the United States, including London (B.1.1.7), Brazil (P.1.), South African (B.1.351), and California strains (B.1.429 and B.1.427), as variants of concern (VOC); these more transmissible, virulent strains are becoming dominant within populations rapidly. VOCs have accumulated key mutations, particularly in the spike protein within the NTD and the RBD, and significant concerns are developing around the efficacy of currently available treatments and vaccines. Understanding factors that underly the level of defense provided by SARS-CoV-2 vaccines against the reference WIV04 strain and the circulating variants of concern is an urgent priority^15^.

We previously reported that seroprevalence among a cohort of uniformly exposed emergency department health care providers was about 5% after the first peak of the infection in Washington, DC, in the late Spring of 2020^16^. PCR confirmed documented SARS-CoV-2 infection increased to about 11% in this cohort of 237 participants by the end of 2020, an incidence of infection in line with health care providers generally in North America^17^. Here we investigate the longitudinal antibody response to the reference Wuhan strain (WIV04), the emerging variants of concerns, and the four HCoVs at three-time points: the Spring of 2020 after the pandemic’s first peak; January of 2021 in the period immediately following the roll-out of the mRNA vaccines; and after the majority of the cohort had been fully vaccinated in early March 2021. By combining PCR and serology test results, we first determined that the previous exposure to SARS-CoV-2 irrespective of the symptoms within this cohort is about 14% before vaccination. We then evaluated SARS-CoV-2 mRNA vaccine-induced antibody isotypes, the magnitude of spike-specific and cross-reactive antibodies, and neutralizing antibodies against the reference WIV04 strain and the circulating VOCs.

## Results

### Determination of SARS-CoV-2 serostatus among mRNA vaccine recipients

Between June 2020 and March 2021, we enrolled 237 healthcare workers in a large tertiary academic medical center (George Washington University, United States), to estimate baseline serostatus and study antibody response after doses 1 and 2 of an mRNA vaccine among previously infected and naïve individuals (Supplementary Figure 1 and table 1). Overall, participants were young (median age=30 years) and healthy (82.1% reported no chronic medical conditions). Of 237 healthcare workers, 161 participants received the Pfizer-BioNTech mRNA vaccine (BNT162b2), and 7 participants received then Moderna mRNA-1273 vaccine. Both of these vaccines use mRNA to induce the expression of stabilized fulllength SARS-CoV-2 spike protein^18^. We collected a total of 424 longitudinal samples from the 237 health care workers. This was done pre-vaccination (n=136 collected 6 months before vaccination), after dose 1 (n=149 collected between 6 – 28 days of vaccination [median 21 days]), and after dose 2 (n=139 collected within 66 days of vaccination [median 54 days]). At the start of the enrollment in June 2020, 7 participants were seropositive, and between June 2020 and Dec 2020, 19 additional participants experienced symptomatic, mild SARS-CoV-2 infection, detected by RT-PCR. None of the infected HCP in this study experienced severe disease requiring hospitalization. To determine the SARS-CoV-2 serostatus of all the participants, we measured SARS-CoV-2 nucleocapsid antibodies in pre-vaccine and dose 1 samples and Spike RBD antibodies before vaccination (Supplementary Figure 1). We stratified the 237 participants as 35 seropositive and 202 as seronegative by combining RT-PCR and serology test results (Supplementary Figure 1). Among the 168 mRNA vaccine recipients, 20 were previously infected, and 148 were naïve. Analysis of self-reported symptoms following vaccination indicated that naïve individuals tended to experience fewer symptoms following both vaccine doses than previously infected individuals (Supplementary Figures 2–3).

### High levels of spike IgG over IgM and IgA antibodies after dose 1 and dose 2

In symptomatic SARS-CoV-2 infections, IgG, IgM, and IgA antibodies are typically developed after 9 days post symptom onset^2^. To understand the antibody response following the doses 1 and 2 mRNA vaccines, we measured the SARS-CoV-2 antibody isotypes against full spike and RBD antigens (Figure 1). Among the naïve, antibody levels were low up to 7 days, and IgG antibodies to full spike and RBD antigens seroconverted in >95% of the participants by day 8 (Figures 1A and D). The predominant response was for the IgG antibodies in both naïve and previously infected participants after doses 1 and 2. Among naïve individuals, IgM and IgA antibodies were higher after the first dose than the second dose (Figure 1G and H). In previously infected individuals, IgM levels were generally lower than the naïve individuals. However, IgA levels in the previously infected individuals were comparable to the naïve individuals after dose 1, and was unchanged after dose 2 (Figure 1H).

**Figure 1.**
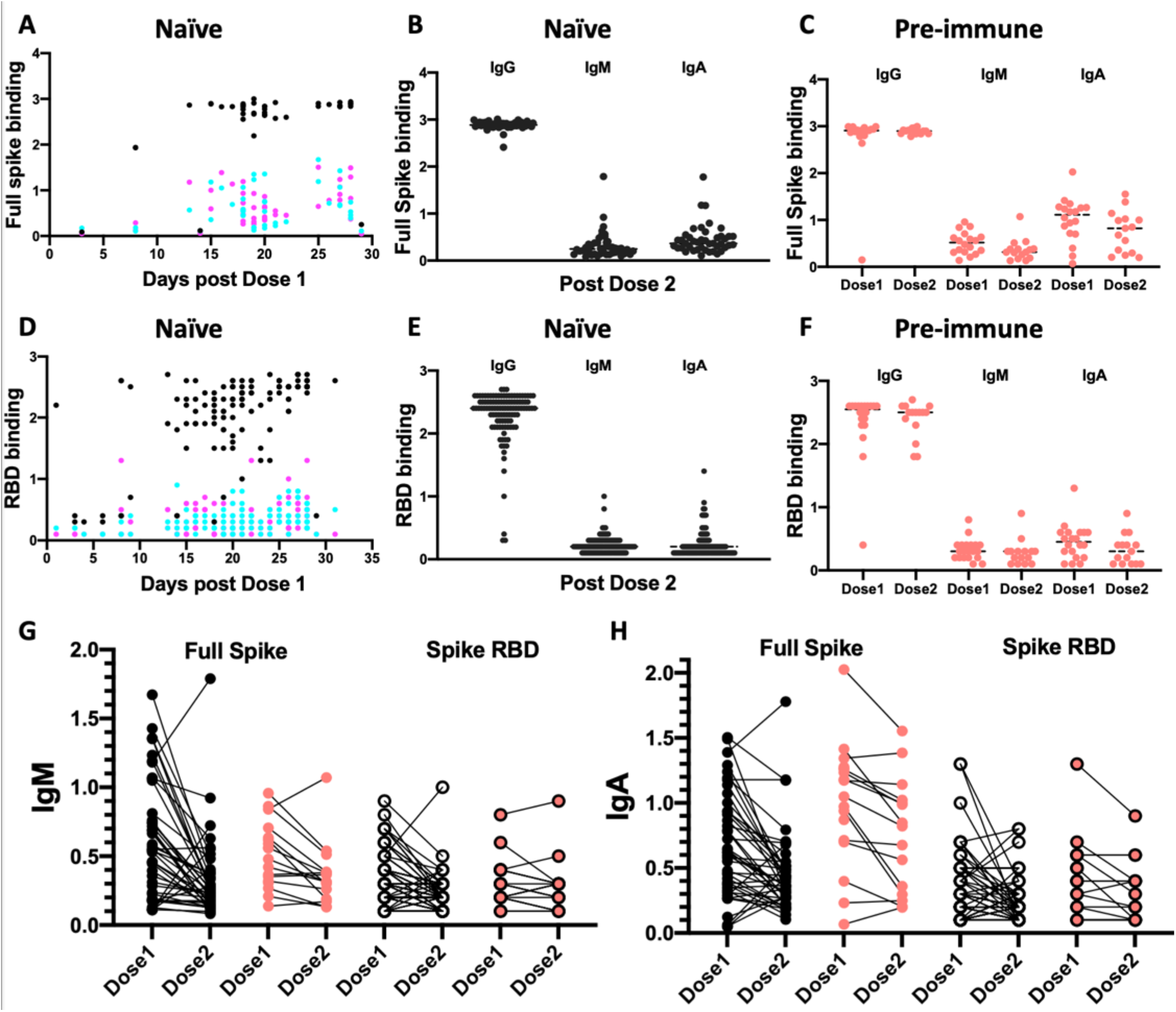
Antibody isotype profiling after mRNA vaccination in naïve and previously infected adults. Longitudinal analysis of serum binding antibodies to (A) full spike (D) RBD after dose 1 in naïve participants. IgG, black; IgM, pink; IgA Cyan. Levels of IgG, IgM and IgA antibodies binding to (B) full spike (E) RBD after dose 2 in naïve participants. Levels of IgG, IgM, and IgA antibodies binding to (C) full spike (F) RBD after dose 1 and 2 vaccination in previously infected participants. Changes in (G) IgM and (H) IgA antibody levels between dose 1 and 2 vaccination in naïve and previously infected participants. ELISA binding is shown as optical density units at 450 nm.

### Comparable spike IgG levels after Dose 2 in previously infected and naïve individuals

Having observed that the IgG antibodies are dominant, we compared the magnitude of IgG response after doses 1 and 2 in titration ELISA assays against full spike, RBD, and NTD antigens (Figure 2). The spike IgG antibody levels, measured by the area under the curve in titration experiments, was robustly boosted among the previously infected participants following dose 1, but this response varied in some individuals (Figure 2A–C). The variation of the spike IgG antibodies after the first dose was higher in naïve than the previously infected individuals. After dose 2, IgG response to spike antigens was highly focused, and the antibody levels were comparable between naïve and the previously infected individuals. The magnitude of RBD antibodies correlated well to NTD and the full spike antibodies in both groups (Figure 2E and F).

**Figure 2.**
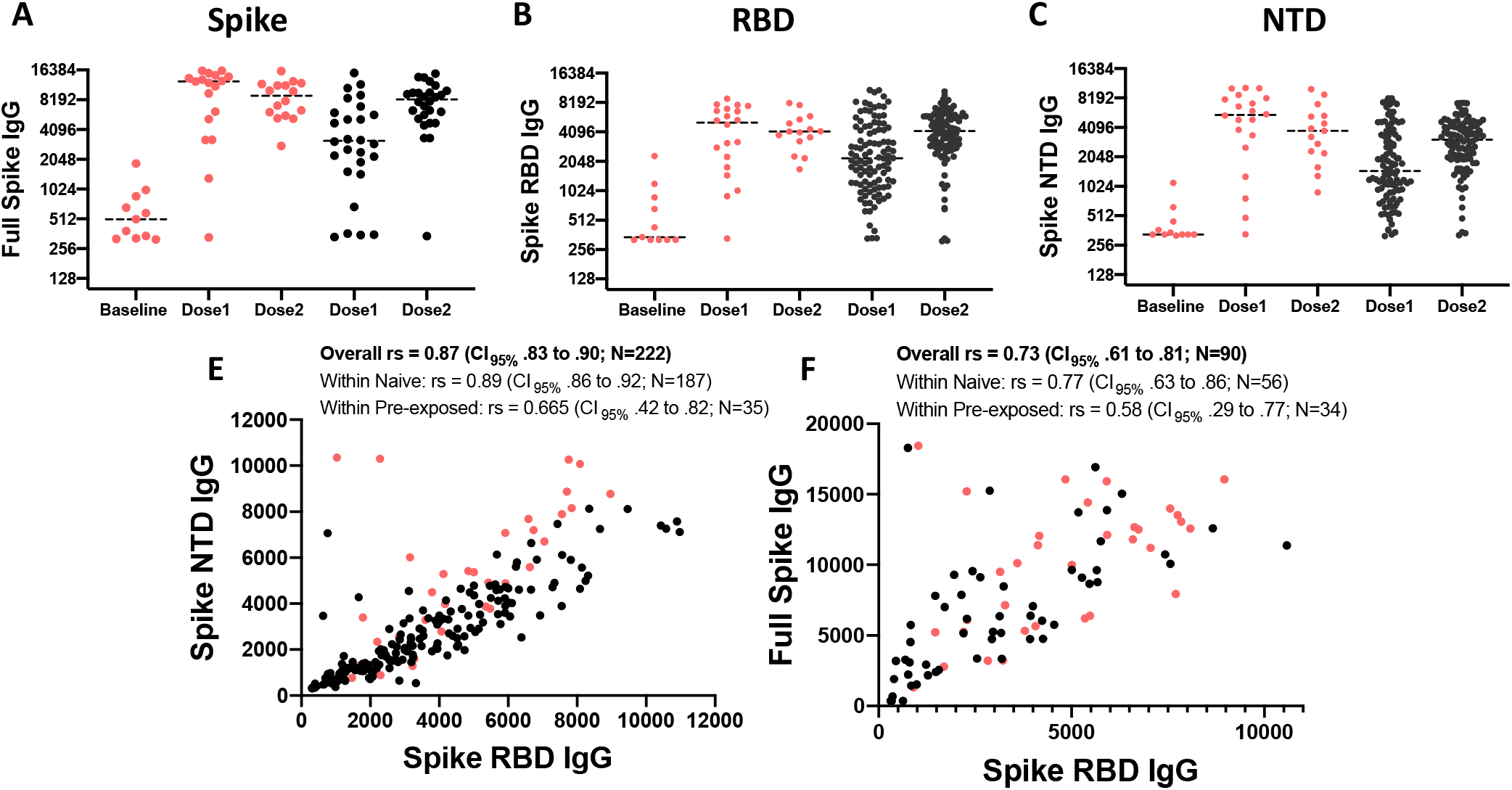
Analysis of spike IgG antibodies after mRNA vaccination in naïve and previously infected adults. Binding IgG levels to (A) full spike, (B) RBD, and (C) NTD antigens were measured by ELISA using serially diluted sera from naïve and previously infected participants after the first and second doses of mRNA vaccination. E) Correlation of binding IgG antibodies between Spike NTD and Spike RBD (F) Correlation of binding IgG antibodies between Spike RBD and full-Spike. Binding antibody titers are expressed as area-under-the-curve (AUC). The nonparametric Spearman correlation coefficient (rs) and the associated 95% confidence interval is shown for previously infected and naïve participants. Black dots, Naïve participants; Red dots, previously infected participants.

### Naïve individuals develop weaker neutralizing antibodies than previously infected individuals

We and others have previously shown that the levels of RBD binding antibodies correlated to the SARS-CoV-2 neutralizing antibody titers^19–22^. To understand the relationship between neutralizing antibodies and spike binding antibodies after mRNA vaccination, we measured live virus SARS-Cov-2 neutralizing titers in 37 paired samples comprising 15 previously infected and 22 naïve individuals after vaccine doses 1 and 2 (Figure 3). Among previously infected and naïve vaccine recipients, we observed a robust correlation between SARS-CoV-2 neutralizing antibodies and the levels of the spike RBD and NTD IgG binding antibodies and a modest correlation with full-spike binding IgG antibodies (Figure 3A and B). We also noticed a moderate-to-high correlation between the neutralizing antibody levels and RBD and full-spike serum IgA antibodies among previously infected vaccine recipients.

**Figure 3.**
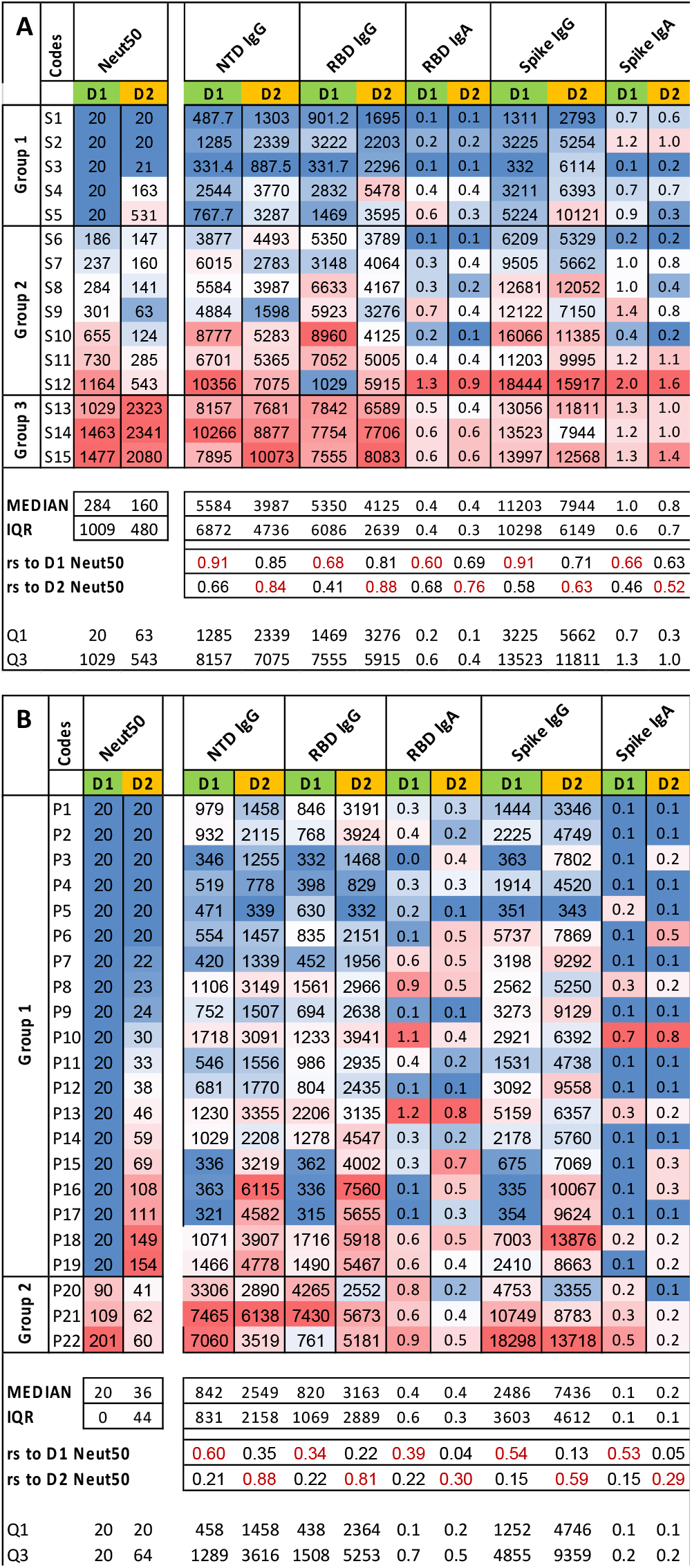
Analysis of SARS-CoV-2 live virus neutralizing antibody titers after mRNA vaccination in naïve and previously infected adults. (A) Heatmap showing the comparison of the neutralizing antibody titers to spike binding antibodies in (A) previously infected (B) naïve participants after the first and second doses of mRNA vaccine. Median, interquartile range (IQR), and the nonparametric Spearman correlation coefficient (rs) are shown for each group. Group stratification by neutralizing antibodies (Group 1, undetectable after dose 1 and remained undetectable or became detectable after dose 2; Group 2, declined between doses 1 and 2; Group 3, improved between doses 1 and 2) are shown.

Strikingly, about 80% (19/22) of the naïve did not develop detectable levels of neutralizing antibodies after dose 1, and about 50% (9/19) of those naive vaccine recipients neutralizing antibodies was not detectable even after dose 2 (Figure 3B). In contrast, 66% (10/15) of previously infected individuals developed neutralizing antibodies after dose 1, and after dose 2, 80% (12/15) of the previously infected had neutralizing antibodies (Figure 3A). Even though both naive and previously infected individuals developed similar levels of spike binding antibodies after dose 2, the mean and median neutralizing antibody levels among the naive recipients were at least ten and four folds weaker than the previously infected vaccine recipients (Cf. Figure 3A and 3B). Overall, the relationship between spike binding and SARS-CoV-2 neutralizing antibodies typically improved between doses 1 and 2, indicated by the increase in correlation and decreased interquartile range (IQR), and followed nonmonotonic relationships, which we divided into three groups. In group 1, neutralizing antibodies were undetectable after dose 1 and remained undetectable or became detectable after dose 2. In group 2, neutralizing antibody response declined between doses 1 and 2, whereas neutralizing antibody response in group 3 improved between doses 1 and 2.

### Neutralizing antibodies developed against WIV04 strain are weaker against other circulating variants

To understand the vaccine effectiveness against the circulating variants (B.1.1.7, B.1.351, and P.1), we analyzed and compared the neutralizing activity for the reference WIV04 strain and the variants after doses 1 and 2 among naive and previously infected individuals in a multiplex surrogate neutralization assay (Figure 4A and B). The multiplex surrogate neutralization assay simultaneously measured antibodies that can block the interaction between RBD and ACE2 in a panel of spike antigens from the reference WIV04 strain and the three most concerning novel viral variants B.1.1.7, P.1, and B.1.351. The percentage of the ACE2 blocking antibodies in the surrogate neutralization assay with 100X diluted sera robustly correlated to the neutralizing antibody titers obtained from the BSL-3 SARS-CoV-2 neutralization assay (Figure 4C). In general, previously infected vaccine recipients displayed higher spike-ACE2 blocking activity compared to the naïve vaccine recipients. The magnitude of RBD binding antibodies correlated with the spike-ACE2 blocking activity against the reference strain and followed a non-monotonic relationship across the three groups as observed for the live virus neutralization assay (Figure 3). The spike-ACE2 blocking activity of B.1.1.7 strains in naïve and preexposed individuals was slightly lower but closely tracked the ACE2 blocking activity of WIV04 (Figure 4D and E). In comparison, the naïve and previously infected vaccine recipients displayed significantly reduced ACE2 blocking activity against B.1.351 and P.1 strains. Overall, people with high RBD binding antibodies also developed better ACE2 blocking activity and vice versa (Figure 4D and E).

**Figure 4.**
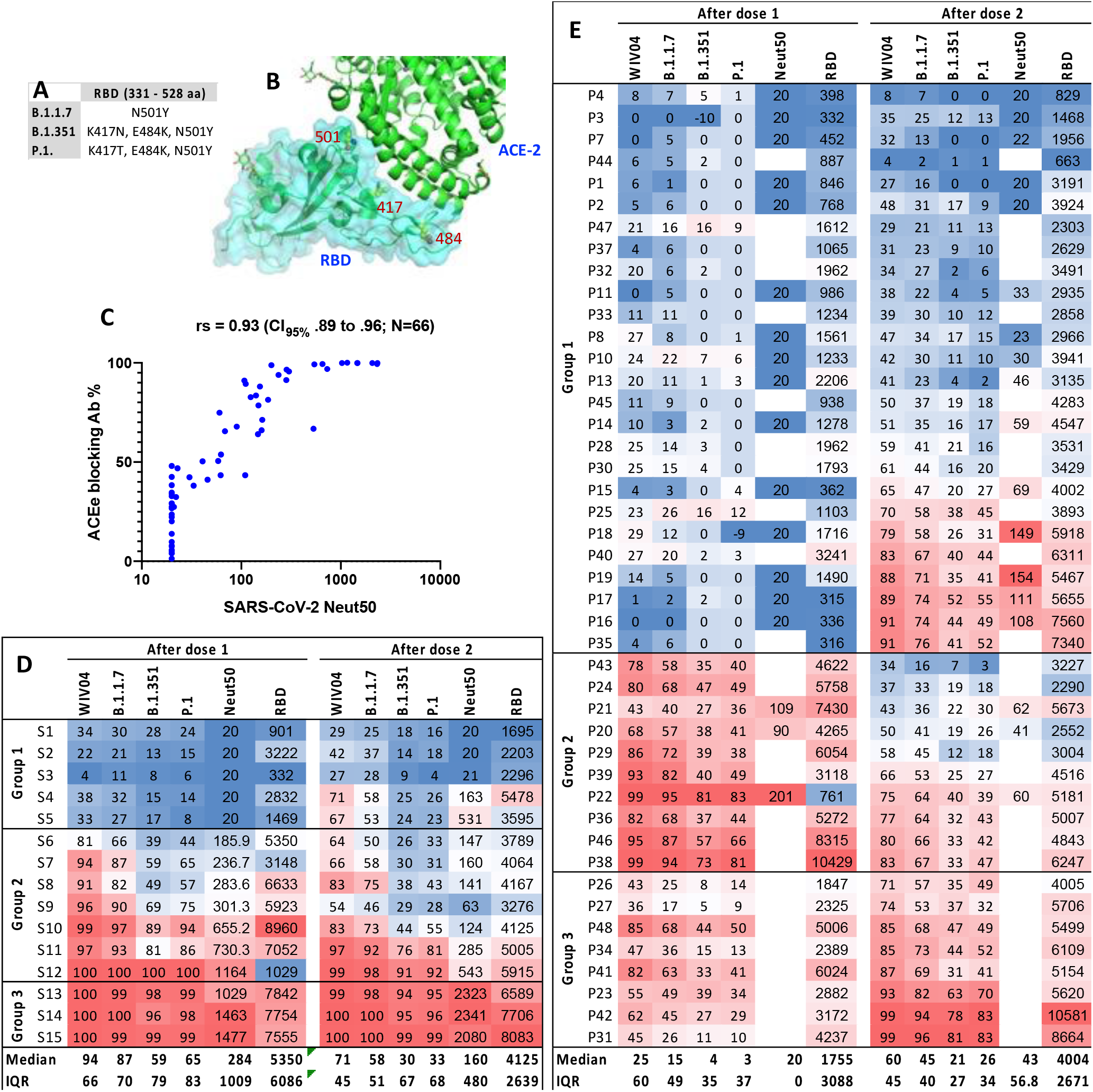
Analysis of mRNA vaccine-induced neutralizing antibody response against circulating variants of concern. (A) The three most concerning circulating variants with single amino acid substitutions within RBD. (B) Structure of RBD complexed with its receptor ACE-2. Amino acid position undergone substitution is shown (PDB ID: 6VW1). (C) Correlation between ACE-2-RBD blocking antibodies and SARS-CoV-2 live virus neutralization titer. The nonparametric Spearman correlation coefficient (rs) and 95% confidence interval are shown. Heatmap showing the comparison of ACE-2 blocking activities against referent and the three variants in (D) previously infected (E) naïve participants after the first and second doses of mRNA vaccine. RBD IgG antibody binding titers (AUC) and live virus neutralization titers (Neut50) are shown for comparison. Median and interquartile range (IQR) are shown. Group stratification by neutralizing antibodies (Group 1, undetectable after dose 1 and remained undetectable or became detectable after dose 2; Group 2, declined between doses 1 and 2; Group 3, improved between doses 1 and 2) are shown.

### mRNA vaccine induces higher levels of endemic HCoV cross-reactive antibodies in previously infected than in naïve individuals

All participants in our cohort have been previously exposed to more than one human endemic CoVs. The development of cross-reactive antibodies to human endemic CoVs was reported previously in hospitalized patients with severe SARS-CoV-2 symptoms^23^. We, therefore, measured longitudinal antibody levels against the full spike antigens from the reference SARS-CoV-2 and the four-human endemic CoVs using a titration ELISA with the samples collected at pre-vaccination and after doses 1 and 2 from previously infected and naïve individuals (Figure 5). While most vaccine recipients developed antibodies to SARS-CoV-2 as expected, we observed that some previously infected and naïve individuals developed strong cross-reactive antibodies to HCoV spike antigens. The cross-reactive antibody levels were more robust after dose 1 against the β-HCoVs (OC43 and HKU-1, Figure 5B–C, G–H and K) than α-HCoVs (NL63 and 229E, Figure 5D–E, I–J and K). Similarly, the cross-reactive antibody levels against HCoVs were more pronounced in previously infected individuals than the naïve individuals, marked by a sharper rise after dose 1 followed by a noticeable decline between the first and second dose (Figure 5). Notably, the levels of cross-reactive antibodies induced after vaccination was highly correlated among HCoVs, indicating that the crossreactive antibodies are most likely targeting the conserved S2 segment of the spike protein (Figure 5L).

**Figure 5.**
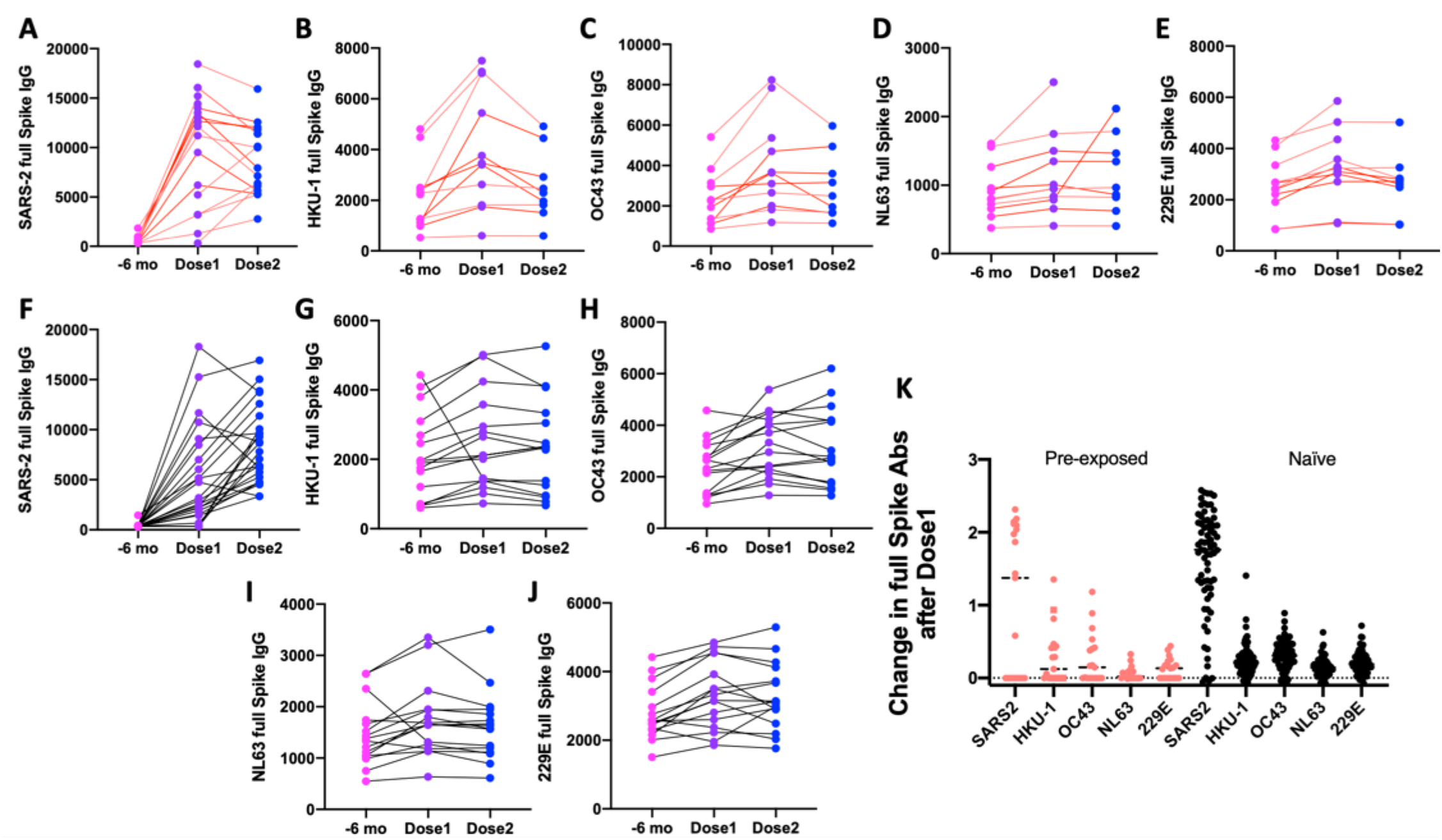
Longitudinal analysis of cross-reactive antibody response against human endemic CoVs after mRNA vaccination. Analysis of IgG binding levels to full spike protein from (A, F) SARS-CoV-2, (B, G) HKU-1, (C, H) OC43, (D, I) NL63, and (E, J) 229E at pre-vaccine, after the first and second doses of mRNA vaccination. IgG antibodies to full spike antigens were measured by ELISA using serially diluted sera from naïve (black line) and previously infected (red line) participants. (K) Analysis of change in spike antibody levels between pre-vaccine and after vaccination in previously infected and naïve participants.

## Discussion

Our study on binding and neutralizing antibody response after two doses of one of the two available mRNA vaccines in 168 naïve and previously infected individuals provides novel insights into vaccine-induced immunity and protection against reinfection by both the reference strain and the circulating variants of concern. Here we report on a lower-than-expected frequency of neutralizing antibodies among fully vaccinated naïve recipients and also provide evidence that some previously infected people may require more than one mRNA vaccine dose to strengthen the immune response against SARS-CoV-2.

As has also been reported in a few independent studies^24,25^, we observed comparable and robust IgG responses to spike RBD and NTD in both naïve and previously infected people after full immunization with an mRNA vaccine^24,25^. Previously, we and others have shown that levels of RBD antibodies correlated well to the magnitude of the neutralizing antibodies in symptomatic SARS-CoV-2 infections^19–22,26^. Among convalescent plasma donors, who recovered after symptomatic infections, 80% had detectable levels of live virus-neutralizing antibodies^27^; of those with binding anti-RBD IgG over 1:160 titers, 95% displayed live virus-neutralization. By contrast, in our cohort, even though binding anti-RBD titers were over these thresholds after full immunization, about 2/5^th^ of the naïve and 1/5^th^ of the previously infected vaccine recipients did not develop detectable levels of neutralizing antibodies as evaluated in an authentic live-virus neutralization assay.

Threshold levels of neutralizing antibodies for SARS-CoV-2 immunity have yet to be established, and even relatively low levels of neutralizing antibodies have been associated with protection in nonhuman primate models^28^. Nonetheless, the lack of detectable levels of neutralizing antibodies in 40% of naïve participants is concerning and differs from the previous reports^25^. That being said, and while data is scanty, clinical evidence does not yet exist that vaccination of the previously infected confers more immunity to reinfection than does vaccination of naïve recipients to primary infection^29^. Moreover, while elevated levels of neutralizing antibodies are considered highly protective^30^, some reports have questioned this relationship^31^, and suggested that other biomarkers may better predict SARS-CoV-2 immunity^32–36^.

Along with neutralizing IgG, the production of anti-spike IgA in the airways has been proposed to prevent SARS-CoV-2-infection^37,38^. Mucosal immunity plays a vital role in viral respiratory infections; IgA dominates early immune response with development of mucosally oriented plasmablasts in natural SARS-CoV-2 infection^39^. Neutralizing mucosal IgA has been associated with milder SARS-COV-2, while circulating neutralizing IgG with more severe disease^34^. The importance of mucosal protection in SARS-COV-2 has led some to advocate for nasally administered vaccines that induce more robust mucosal IgA responses^40^. In our study parenterally delivered mRNA vaccines induce high levels of anti-Spike IgG—similar to what is seen in severe disease. IgA production, by contrast, is induced in more modest quantities, comparable to what is seen in mild disease^11^. Induction of strong IgG and, low IgA levels in naïve patients after receiving one of the two mRNA vaccines has also been recently reported in other independent studies^42^. Serum IgA levels are not a direct measure of secretory IgA levels, since systemic IgA are predominantly made of monomeric IgA1 subclass, and mucosal IgA is a polymeric IgA2 subclass^43^. However, induction of mucosal IgA after parenteral mRNA vaccination has been reported^41^. Our data shows that after full vaccination, spike IgA levels in serum correlated better to the levels of neutralizing antibodies in previously infected vaccine recipients than among naïve recipients, a phenomenon that may be related to a more mature immune response in the former group and recall of memory B cells from natural infection.

Multiple reports have surfaced recently that previously infected individuals may not need or benefit from a second dose of an mRNA vaccine^24,44–48^. Our data, by contrast, shows that 1/3^rd^ of previously infected individuals did not produce detectable levels of neutralizing antibodies after the first dose (Figure 3A, Group 1); after the second dose, however, 40% of these non-responders produced neutralizing antibodies. A lack of increased spike binding antibody level after the second dose in those previously infected with SARS-CoV-2 has also been cited as evidence that the second dose is unnecessary^24,49,50^. Our data also shows a stable or declining antibody levels after the second dose in 2/3^rd^ of previously infected individuals (Figure 3A, group 2 or 3), a result that has also been noted by other^49,51,52^. Binding antibody levels after the first dose in the previously infected tend to increase by orders of magnitude; what our data shows is that in this group while after the second dose of vaccine higher antibody levels tend to decline, lower levels tend to increase, suggesting focusing of the immune response. This observation is strengthened by the fact that the proportion of previously infected subjects with neutralizing antibodies increased even while mean levels of binding antibodies stayed level or decreased, likely an effect of memory B cells, and associated plasma B cells in the previously infected producing more targeted and affinity-matured antibodies^17,53^.

Compared to naïve recipients, which experience significant increases in the production of variant-neutralizing antibodies after both doses of vaccine, we observed a trend toward decreasing neutralization against the circulating SARS-CoV-2 variants in the previously infected after the second dose. This may be related to immune dominance of ‘originally imprinted’ antigens that can then weaken the response to subsequently encountered related antigens, as could be the case with the SARS-CoV-2 variants^54,55^. Others have argued that ‘affinity maturation’ can over time produce ‘broadly neutralizing antibodies’ (BNABs) with the ability to provide sterilizing immunity based on high affinity for conserved epitopes among variant pathogens^53,56^. ‘Persistent’ exposure to pathogen antigens is thought to underlie the process of affinity maturation and the development of BNABs. While optimized^53^ timing of SARS-CoV-2 spike antigen exposure through booster vaccinations with the currently approved mRNA vaccines may be a route to increase resistance to SARS-CoV-2 variants, our data doesn’t suggest this. The effectiveness of the neutralizing antibodies developed against B.1.1.7 and B.1.351 variants in those who received a BNT162b2 mRNA vaccine, have been reported as 90% and 75%, respectively^57^. There are reports^58^ of the development of adequate neutralizing antibodies against SARS-CoV-2 variants following administration of the existing mRNA vaccines to previously infected subjects. However, our data suggests that as the immune response to SARS-CoV-2 spike antigen matures in vaccinated subjects, susceptibility to the variants will remain significant, raising the question if variant-focused re-vaccination with modified vaccines will become necessary? There are concerns^55,59^ regarding potential problems with the effectiveness of such modified vaccines; however, there are not at present, strong reasons to think that the current vaccine regimen, or repeated doses of the existing vaccines, are likely to induce BNABs against SARS-CoV-2 variants. By contrast, a potent cross-reactive antibody^60^ derived from a SARS-CoV-2-variant epitope has recently been reported, suggesting that modified vaccines produced in this fashion may prove more broadly effective. Widely administered booster immunizations against reference/variants will expose a previously vaccinated and sensitized population to risk of vaccine reaction with attendant risks and discomforts^61^. Our data shows that previous SARS-CoV-2 exposure either from infection or vaccine results in greater symptoms in response to a second exposure to SARS-CoV-2 antigens; however, a recent small scale study which looked at booster vaccinations with one of the two mRNA vaccines reported no more than moderate symptoms in just 15% of subjects^62^.

In our study naïve subjects experienced significantly increased spike antibody levels against the SARS-CoV-2 reference strain, SARS-CoV-2 variants, and the human endemic HCoVs after both doses of vaccine. Previously infected subjects after dose 2, by contrast, saw modest improvement in neutralizing antibodies against the Wuhan strain, but antibody levels were stable or declining against HCoV, and SARS-CoV-2 variants. HCoV memory B and T cells have been implicated in the early response to SARS-CoV-2 exposure^17^ and likely explain the induction of HCoV antibodies in infection/vaccination after the first dose in both the previously infected and the naïve. As has been shown with Zika and Dengue viruses^63,64^, which are closely related flaviviruses, transient cross-reactive antibodies develop in the acute phase^37^ and tend to fade as immune responses mature. Nonetheless, cross-reactive HCoV antibodies targeting the S2 segment were reported to be boosted in some hospitalized patients^13,23,65^ and have been associated with the development of neutralizing Ab^37^.

Reliable serological assays to monitor community transmission of SARS-CoV-2 and its variants, and to support post-acute SARS-COV-2 (PASC) studies are urgently needed. After vaccination, spike antigen serology is unsuitable for detecting SARS-CoV-2 infection; in those immunized with one of the currently available mRNA vaccines, or similar, which do not expose the recipient to the totality of viral antigens, serological assays targeting antigens not included in the vaccine may be used to identify evidence of previous infection. Currently antibodies against the nucleocapsid phosphoprotein are used for this purpose, however, our analysis of longitudinal samples shows that nucleocapsid serology displays poor sensitivity over time (Supplementary Figure 1E). SARS-CoV-2 non-structural proteins such as open reading frame ORF-8 and ORF-3b^66,67^ have been identified as alternate antibody targets, which in combination may improve sensitivity for detecting previous SARS-CoV-2 infection. These and other non-spike viral epitopes^68^ constitute a promising strategy to distinguish past SARS-CoV-2 infection from vaccination.

## Conclusion

Our data shows a significant lack of neutralizing antibodies in naïve subjects after full vaccination, and a more limited but concerning lack of neutralizing antibodies in some previously infected individuals. In contrast to what has previously been reported, we find that previously infected individuals may benefit from two doses of the currently available mRNA vaccines. Our data does not show improving neutralizing antibodies against circulating variants with repeated vaccine doses in the previously infected, suggesting that booster vaccinations with vaccines modified for the variants may be called for in the future. We also find that nucleocapsid IgG wanes over time and is therefore limited as an assay for previous infection with SARS-CoV-2.

Study limitations include the inclusion of subjects receiving only mRNA vaccines. Our cohort’s previously exposed study subjects were unlikely to have been infected with one of the variants. Study participants included primarily young, healthy adults; chronically ill, immunocompromised, and the very young and very old are not represented in our study. Our data represent interim values that are likely to evolve with time in terms of response to vaccination.

## Experimental Methods

### Clinical study

In this study, a total of 424 venous blood samples were collected from 237 unique ED HCP participants at George Washington University Hospital (GWUH) during the timeframe of June 2020 to March 2021. ED HCP participants were defined as any GWUH ED healthcare providers that came into close contact with patients between December 2019 and March 2021. Clinical roles of study participants included physicians, advanced practice providers, nurses, technicians, respiratory technicians, and environmental services personnel. Samples were collected during three separate time periods. Testing occurred in May/June 2020, January of 2021, and March 2021. Participants were encouraged to participate in subsequent testing rounds so that interval serologic changes in response to SARS-CoV-2 exposures or vaccinations could be analyzed. The study was approved by the George Washington University IRB#: NCR202406.

#### Patient Recruitment

Emails were sent to all GWUH ED HCP personnel through an ED staff listserv. Additionally, in order to reach those not on the listserv, notifications were sent out through GWUH’s ED nurse/technician scheduling system and fliers were placed in break rooms with QR codes connecting to patient sign-up forms. Participation days and times overlapped nurse and technician shift changes in order to encourage on-going and off-going staff to participate. All ED HCP personnel who chose to participate in this study provided written informed consent. All personnel who consented to participate were included in the study.

#### Data Collection

All participants were asked to complete a questionnaire about demographics (age, gender, race/ethnicity, home city), ED HCP occupation, non-ED HCP affiliations (e.g. also work in the ICU), past medical history (PMH), current medications, smoking history, history of known positive COVID-19 status, recent/intercurrent viral syndrome symptoms (fever, fatigue, dry cough, anorexia, body aches, dyspnea, sputum, sore throat, diarrhea, nausea, dizziness, headache, vomiting, and abdominal pain), relative number of COVID-19 exposures at work and outside of work, and personal protective equipment (PPE) wearing habits (time spent wearing and frequency of changing surgical masks, N95 masks, and Powered Air Purifying Respirators. Additionally, participants in the second and third rounds of testing were asked questions regarding COVID-19 vaccine status, type of COVID-19 vaccine received, dates of first and second vaccine doses, and recent/intercurrent viral syndrome symptoms after obtaining COVID-19 vaccinations.

#### Sample Collection

Venous blood samples were collected from each participant during each round of testing. 10 ML of blood was drawn into an SST tube, and refrigerated overnight to allow for serum separation. Subsequently the serum was drawn off into a 2 ML Eppendorf tube and stored at −80 deg. C. for subsequent use.

#### Protein expression and purification

The expression and production of halo-tagged SARS-CoV-2 and the four human endemic CoV RBD antigens from mammalian cells were previously described^22^. The halo-tagged SARS-CoV-2 NTD antigen (16–305 amino acids, Accession: P0DTC2.1) was designed and expressed in mammalian Expi293 cells as described for RBD antigens. RBD and NTD antigens were site-specifically biotinylated using Halotag PEG biotin ligand (Promega), following the manufacturer’s protocol. For producing SARS-CoV-2 spike protein trimer, a codon-optimized synthetic gene was synthesized to encode for a prefusion-stabilized SARS-CoV-2 spike protein (16-1208 amino acids, Accession: P0DTC2.1) with an N-terminal human serum albumin secretion signal peptide and C-terminal T4 foldon trimerization domain, TEV protease cleavage site, His8 tag, and the twin-strep tag. The prefusion-stabilized SARS-CoV-2 construct contains two consecutive proline substitutions in the S2 subunit as described before (PMID: 32075877, PMID: 32155444). The synthetic gene was cloned between KpnI and XhoI sites of the mammalian expression plasmid pαH. SARS-CoV-2 spike protein was expressed as described for RBD and NTD antigens in Expi293 cells and purified from mammalian cell culture supernatant using Strep-Tactin immobilized affinity resin (IBA Lifesciences). The bacterial expression construct for full-length SARS-CoV-2 nucleocapsid was a gift from Nicolas Fawzi (Addgene plasmid # 157867; http://n2t.net/addgene:157867; RRID:Addgene_157867)^69^. The MBP fused nucleocapsid protein was expressed and purified from BL21(DE3)PlysS as described before^70^. The purified full-length ectodomain of the human coronavirus spike proteins (HCoV-NL63, 40604-V08B; HCoV-OC43, 40607-V08B, HCoV-229E, 40605-V08B, and HCoV-HKU1, 40606-V08B) were purchased from Sino Biological.

#### Generation 2 RBD or NTD ELISA

All serum samples tested by ELISA assay were heat-inactivated at 56°C for 30 min to reduce risk from possible residual virus in serum. Briefly, 50 μL of Streptavidin (Invitrogen™ cat # 434302) at 4 μg/mL in Tris-Buffered Saline (TBS) pH 7.4 was coated in the 96-well, high-binding microtiter assay plate (Greiner Bio-One cat # 655061) for 1 hour at 37°C. The coating solution was removed, then 100 μL of blocking solution, 1:1 NAP-Blocker (NAP [Non Animal Protein]-BLOCKER™, G-Biosciences cat # 786-190) in TBS was added for 1 hour at 37°C. Serum samples were diluted at 1:40, or serially diluted (1:100 – 1:8100), in 3% Bovine Serum Albumin (BSA) in TBS containing 0.05% Tween 20 (TBST) with biotinylated spike RBD or NTD antigen at 1 μg/mL in a 96-round-well V bottom plate (Diaago cat # R96-300V) and incubated for 1 hour at 37°C. The blocking solution was removed, then 50 μL of diluted serum was added to the assay plate and incubated for 15 minutes at 37°C. The plate was washed three times using wash buffer (TBS containing 0.2% Tween 20), then 50 μL of horseradish peroxidase-conjugated secondary Goat Anti-Human secondary antibody at 1:40,000 dilution in 3% milk was added for 40 minutes at 37°C. For measuring total Ig, a mixture of anti-IgG, anti-IgM, and anti-IgA were added together (Cat #109-035-008, 109-035-043, 109-035-011, Jackson ImmunoResearch). For measuring isotype specific antibody, only the respective goat antihuman HRP conjugated IgG, IgM, or IgA was used. The plate was washed three times with wash buffer, and 50 μL of 3,3’,5,5’-Tetramethylbenzidine (TMB) Liquid Substrate (Sigma-Aldrich cat # T0440) was added to the plate, and absorbance was measured at 450 nm using a plate reader after stopping the reaction with 50 μl of 1 N HCl (Biotek Epok, Model # 3296573).

#### Full-length Spike Protein ELISA

Briefly, 50 μL of full-length spike protein at 2 μg/mL in Tris-Buffered Saline (TBS) pH 7.4 was coated in the 96-well, high-binding microtiter plate (Greiner Bio-One cat # 655061) for 1 hour at 37°C. The coating solution was removed, then 100 μL of blocking solution (3% milk in TBST), was added for 1 hour at 37°C. Serum samples were diluted at 1:40, or serially diluted (1:100 – 1:8100), in the blocking solution. The blocking solution was removed, then 50 μL of diluted serum was added to the plate and incubated for 1 hour at 37°C. The plate was washed three times using wash buffer (TBS containing 0.2% Tween 20), then 50 μL of horseradish peroxidase-conjugated secondary Goat Anti-Human secondary antibody at 1:40,000 dilution in 3% milk was added for 1 hour at 37°C. For measuring isotype specific antibody, only the respective goat anti-human IgG, IgM, or IgA was used (Cat #109-035-008, 109-035-043, 109-035-011, Jackson ImmunoResearch). The plate was washed three times using wash buffer, then 50 μL of 3,3’,5,5’-Tetramethylbenzidine (TMB) Liquid Substrate (Sigma-Aldrich cat # T0440) was added to the plate, and absorbance was measured at 450 nm using a plate reader after stopping the reaction with 50 μl of 1 N HCl (Biotek Epok, Model # 3296573).

#### Live Virus Neutralization Assay

We utilized a SARS-CoV-2 luciferase reporter virus (nLuc) to assess neutralizing antibody activit^71^. Vero E6 cells were plated at 2×10^4^ cells/well in a black 96-well clear bottom plate (Corning). Heat-inactivated serum was diluted 1:20 initially, followed by a 3-fold dilution series up to eight dilution spots in DMEM supplemented with 5% FBS. Diluted serum was incubated in a 1:1 ratio with SARS-CoV-2-nLuc to result in 75 PFU virus per well. Serum-virus complexes were incubated at 37C with 5% CO_2_ for 1 hour. Following incubation, serum-virus complexes were added to the plated Vero E6 cells and incubated for 48 hours at 37C with 5% CO_2_. After incubation, luciferase activity was measured with the Nano-Glo Luciferase Assay System (Promega) according to the manufacturer specifications. Neutralization titers (EC50) were defined as the dilution at which a 50% reduction in RLU was observed relative to the virus (no antibody) control.

#### Meso-scale multiplex surrogate neutralization assay

A multiplexed Meso Scale Discovery (MSD) immunoassay (MSD, Rockville, MD) was used to measure the ACE-2 blocking antibodies to SARS-CoV-2 reference strain and circulating variants (B.1.1.7, P1, B.1.351) using the MSD V-PLEX SARS-CoV-2 Panel 7 (ACE2) kit according to the manufacturer’s instructions. Briefly, plates were blocked with MSD Blocker A for 30 minutes and then washed three times prior to the addition of reference standard, controls and heat-inactivated samples diluted 1:10 or 1:100 in diluent buffer. Plates were incubated for 1 hour with shaking at 700 rpm. A 0.25μg/ml solution of MSD SULFO-tag conjugated ACE-2 was added and incubated for 1 hour with shaking, plates were washed and read with a MESO QuickPlex SQ 120 instrument. Each plate contained duplicates of a 7-point calibration curve with serial dilution of a reference standard and a blank well. Results were reported as percent inhibition calculated based on the equation ((1 – Average Sample ECL Signal / Average ECL signal of blank well) x 100).

#### Nucleocapsid Assay

Wells of a high-binding microtiter plate (Greiner Bio-One cat # 655061) were coated with 50 μL anti-MBP (New England Biolabs) at 3 ug/mL in Tris Buffered Saline (TBS) pH 7.4, and then blocked with 100 μL of blocking solution (3% non-fat milk in TBS containing 0.05% Tween 20). The plate was washed three times using wash buffer (TBS containing 0.2% Tween 20). 50 μL of 2ug/mL MBP fused full-length nucleocapsid protein and MBP proteins in blocking solution were added to respective wells. The plates were incubated for 1 hour at 37°C. The plates were washed 3 times, then 50 μL of heat-inactivated serum at 1:40 dilution was added to wells containing full-length N protein and MBP proteins were added respectively and further incubated for 1 hour at 37°C. The plates were washed with wash buffer, then 50 μL of alkaline phosphatase-conjugated secondary goat anti-Human anti-IgG (Sigma Cat # A9544), anti-IgA (Abcam Cat # AB97212), and anti-IgM (Sigma Cat # A3437) at 1:2500 dilution was added and incubated for 1 hour at 37°C. The plate was washed, and 50 μL p-Nitrophenyl phosphate substrate (SIGMA FAST, cat N2770) was added to the plate and absorbance measured at 405 nm using a plate reader (Biotek Epok, Model # 3296573). Appropriate control sera were included in the study.

### Statistical analyses

Characteristics of the sample were summarized using simple descriptive statistics. To describe the magnitude and spread of full spike and RBD binding IgG, IgM, and IgA titers, we generated jittered dot plots stratified by antibody isotype, time of sampling (post dose 1 and post dose 2), and prior exposure to SARS-CoV-2. Line graphs stratified by time of sampling were used to describe trends in change in antibody levels between doses. The strength of the correlations between spike RBD and spike NTD titers, between spike RBD and full Spike titers, between spike binding antibody titers and SARS-CoV-2 live virus neutralization titers, and between SARS-CoV-2 live virus neutralization titers and ACE2 blocking Ab percent, were evaluated using a two-tailed Spearman’s rank correlation in Prism 9. Neutralizing antibody responses were grouped into Group 1 (undetectable after dose 1 and remained undetectable or became detectable after dose 2), Group 2 (declined between doses 1 and 2), and Group 3 (improved between doses 1 and 2), and Spearman’s coefficient was estimated to assess correlations between neutralizing and binding antibody titers, stratified by vaccine dose. Crossreactivity between WIV04 and three SARS-CoV-2 variants (B.1.1.7, B.1.351, and P.1), and crossreactivity between SARS-CoV-2 and other endemic human coronaviruses was compared using stratified description of antibody titers and kinetics following each vaccine dose.

## Acknowledgements

This study was supported by grants from the National Cancer Institute (U54CA260543 to A.M.de, R.S.B, and L.P.) and National Institutes of Health, Allergy and Infectious Diseases (U01AI151788 to A.M.de and L.P). We would like to thank the contributions of HCP donors and the staff at GWU for this work. We thank Dr. Liu and the members of the Liu Lab at George Washington University’s Milken Institute School of Public Health. We would like to acknowledge Meso Scale Discovery for donating plates and reagents to perform the surrogate neutralization assay to this study. We would like to thank the Adams School of Dentistry DELTA Translational ReCharge Center and its Co-directors, Drs. Shannon Wallet and Robert Maile for the infrastructure support to this study. We thank Mr. John Forsberg at the UNC protein expression core facility and Mr. Salman Khan for assisting with protein expression.

## Supplementary Information

**Supplementary Figure 1.**
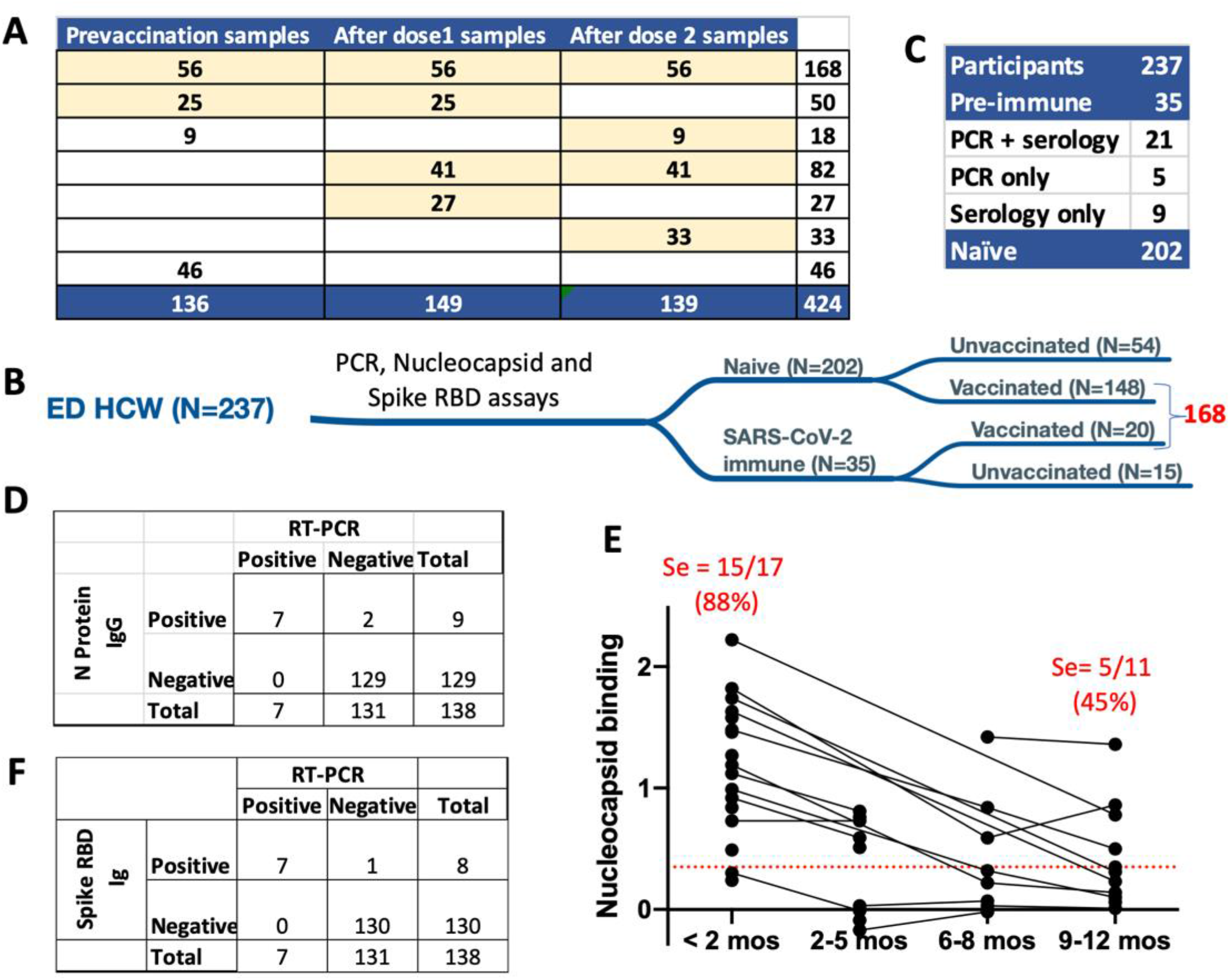
Determination of SARS-CoV-2 serostatus of health care workers. (A) Blood samples stratified by collection timepoints. (B-C) Cohort stratified by SARS-CoV-2 infection and vaccination status. Comparison of SARS-CoV-2 (D) Nucleocapsid IgG and (F) RBD Ig assays performance to RT-PCR test results with prevaccination samples. (E) Nucleocapsid IgG assay sensitivity over time.

**Supplementary Figure 2.**
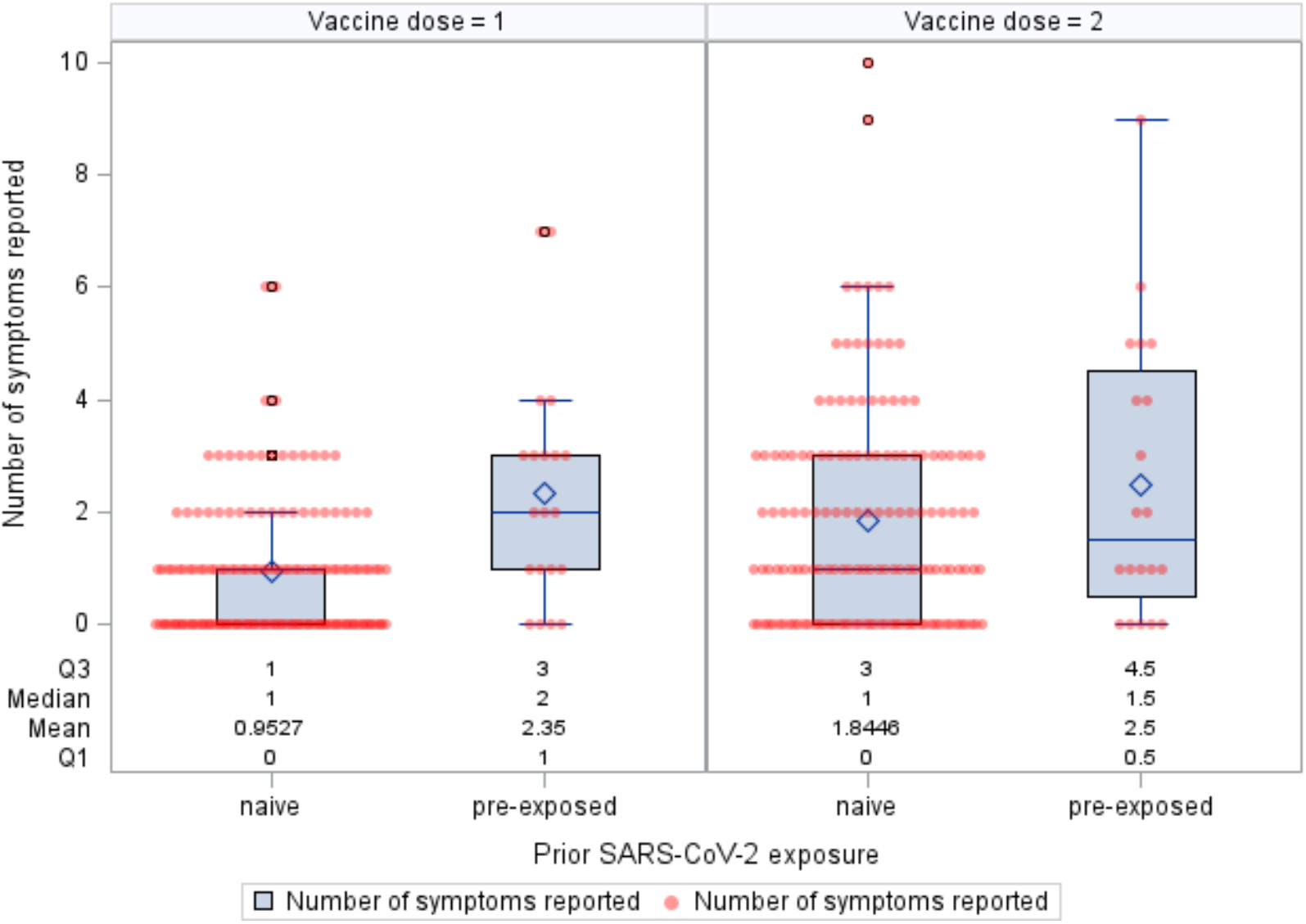
Number of symptoms reported following vaccination, by dose and prior SARS-CoV-2 exposure (n=168). The box plots show the mean, median, and interquartile range for number of symptoms reported following each COVID-19 vaccine dose, stratified by participants who were SARS-CoV-2 naïve and SARS-CoV-2 pre-exposed at pre-vaccination. The red dots represent individual data on number of symptoms reported following each vaccine dose.

**Supplementary Figure 3.**
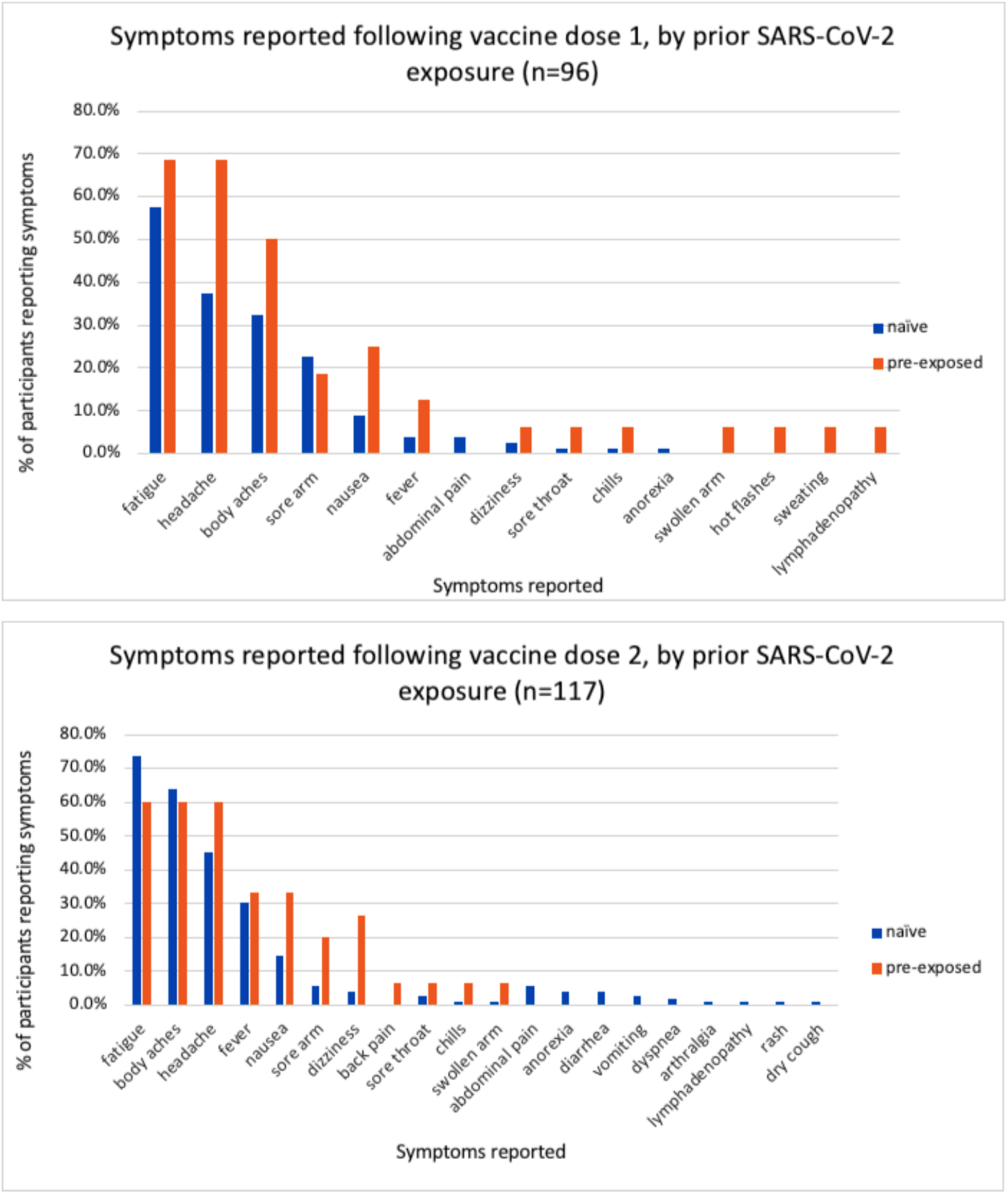
Types of symptoms reported following (A) vaccine dose 1 and (B) vaccine dose 2 stratified by participants who were SARS-CoV-2 naïve and SARS-CoV-2 pre-exposed at baseline.

**Supplementary Table 1.** Baseline characteristics of healthcare personnel who received COVID-19 vaccines.

| Characteristic | All Vaccine Recipients<br>(n=168) |  | Pfizer Vaccine<br>Recipients (n=161) |  |
| --- | --- | --- | --- | --- |
|  | N or Median | % or IQR | N or<br>Median | % or IQR |
| Age (years) | 31 | 26-39 | 31 | 26-37 |
| Sex |  |  |  |  |
| Female | 119 | 70.8 | 117 | 72.7 |
| Male | 49 | 29.2 | 44 | 27.3 |
| Race/Ethnicity (2 missing) |  |  |  |  |
| Asian | 15 | 9.0 | 12 | 7.6 |
| Black or African-American | 14 | 8.3 | 14 | 8.8 |
| Hispanic or Latinx | 5 | 3.0 | 4 | 2.5 |
| White | 126 | 75.9 | 124 | 78.0 |
| More than one race | 6 | 3.6 | 5 | 3.1 |
| Clinical Role (1 missing) |  |  |  |  |
| Attending physician | 35 | 20.8 | 32 | 19.9 |
| Resident or physician assistant | 46 | 27.4 | 44 | 27.3 |
| Nurse | 57 | 33.9 | 56 | 34.8 |
| Respiratory therapist | 1 | 0.6 | 1 | 0.6 |
| Emergency department technician | 28 | 16.7 | 27 | 16.8 |
| Number of chronic medical conditions reported (1 missing) |  |  |  |  |
| 0 | 138 | 82.6 | 132 | 82.5 |
| 1 or 2 | 29 | 17.4 | 28 | 17.5 |
| Smoking behavior (12 missing) |  |  |  |  |
| Non-smoker | 138 | 88.5 | 132 | 88.0 |
| Ever smoker (including current, prior, and vape use) | 18 | 11.5 | 18 | 12.0 |

